# Intranasal L-MYC Neural Stem Cells Restore Neurovascular and Neuroimmune Homeostasis After TBI via CX3CL1 Upregulation and ICAM-1 Downregulation

**DOI:** 10.64898/2026.10.09.756809

**Authors:** Brecken Enright, Jeffrey P. Cheng, Vikram Adhikarla, Isabella Vasquez, Yate-Ching Yuan, Corina O. Bondi, Russell Rockne, Anthony E. Kline, Margarita Gutova

**Affiliations:** Department of Stem Cell Biology and Regenerative Medicine, Beckman Research Institute, City of Hope, 1500 East Duarte Road, Duarte, CA, 91010, USA; Physical Medicine and Rehabilitation, University of Pittsburgh School of Medicine, Pittsburgh, PA, 15224, USA; Safar Center for Resuscitation Research, University of Pittsburgh, Pittsburgh, PA, 15224 USA; Department of Computational and Quantitative Medicine, Beckman Research Institute, City of Hope, 1500 East Duarte Rd, Duarte, CA, 91010, USA; Center for Neuroscience, University of Pittsburgh, Pittsburgh, PA, 15224 USA; Neurobiology, University of Pittsburgh, Pittsburgh, PA, 15224 USA; Center for the Neural Basis of Cognition, University of Pittsburgh, Pittsburgh, PA, 15224 USA; Critical Care Medicine, and Psychology, University of Pittsburgh, Pittsburgh, PA, 15224 USA; Psychology, University of Pittsburgh, Pittsburgh, PA, 15224 USA

**Author notes:** **Corresponding Authors: Margarita Gutova, PhD** Assistant Professor Beckman Research Institute at City of Hope Department of Stem Cell Biology and Regenerative Medicine 1500 E. Duarte Rd. Duarte, CA, 910101 Office: Miller Bld, Room 111 Lab: Kaplan Black Bldg, Room 109B, **Anthony E. Kline**, PhD Professor Physical Medicine & Rehabilitation, Critical Care Medicine, Psychology, and Center for Neuroscience Associate Director of Rehabilitation Research, Safar Center for Resuscitation Research University of Pittsburgh School of Medicine Children’s Hospital of Pittsburgh John G. Rangos Research Center – Room 6126.

**Keywords:** LMNSC neural stem cells, traumatic brain injury, controlled cortical impact brain injury, neurovascular, neuroimmune, CX3CL1, ICAM-1

## Abstract

Traumatic brain injury (TBI) triggers persistent neuroimmune and neurovascular dysfunction that contributes to secondary brain injury and limits recovery. Here, we investigated whether a single intranasal dose of LMNSC01 could reprogram post-traumatic neuroinflammatory and neurovascular responses and whether these effects were conserved across biological sex. Immunocompetent male and female rats underwent a controlled cortical impact (CCI) or Sham surgery and 7 days later received a single bolus dose of LMNSC01 (6 × 10^6^ cells/dose) or vehicle. Integrated histological and transcriptomic analyses and multiplex tissue ELISA were used to characterize injury- and treatment-associated responses.

CCI induced persistent but sex-dependent transcriptional changes. Male rats exhibited prominent innate immune, adhesion, extracellular matrix, and tissue remodeling responses, whereas females demonstrated broader activation of microglial, interferon, complement, MHC/antigen presentation, and humoral/adaptive immune pathways. LMNSC01 treatment reprogrammed these distinct injury states through partially sex-specific mechanisms. In males, LMNSC01 increased endogenous immune-regulatory mediators, including *Il1rn*, and reduced pathways associated with adhesion and immune-cell trafficking. In females, treatment decreased expression of genes related to microglial, leukocyte, MHC/antigen-presentation, cytokine, and adhesion pathways, and reduced macrophage- and CD45-positive cell signatures. Despite these sex-dependent transcriptional changes, LMNSC01 produced conserved protein changes in both sexes, reducing injury-associated ICAM-1 and restoring CX3CL1 proteins, proposing a novel molecular mechanism of LMNSCs modulating endothelial activation and neuron– microglia communication post injury. Our study provides further insight into sex differences in responses to injury and treatment after TBI and highlights candidate repair mechanisms, supporting further development of LMNSC01 as a minimally invasive therapy for TBI.

**Significance Statement:** This study provides evidence that a single intranasal dose of LMNSC01 can reprogram chronic neuroimmune and neurovascular dysfunction after TBI in both sexes. Despite distinct male and female injury and treatment responses, LMNSC01 produced a conserved reduction in ICAM-1 and increase in CX3CL1, suggesting reduced endothelial activation and restoration of homeostatic neuron–microglia signaling. Identification of shared changes in neurovascular and neuroimmune signaling provides a candidate mechanism for neural stem cell-mediated repair and supports the translational development of non-invasive intranasal LMNSC01 therapy for TBI.

## Introduction

Traumatic brain injury (TBI) is a significant health care issue that often results in long-term neurological disabilities and affects approximately sixty-nine million individuals worldwide each year [1, 2]. TBI can negatively affect motor, cognitive, and affective function [3]. Among these behaviors, cognitive impairments, particularly deficits in higher order functioning such as executive function, are often more pronounced and persistent [4, 5] and contribute substantially to reduced quality of life. TBI can also disrupt interpersonal relationships and emotional well-being. The economic cost to society resulting from acute medical and subsequent rehabilitative care along with the loss of productivity due to the inability to return to work accounts for billions of dollars each year [6]. Thus, there is a critical need for effective therapies that can mitigate the biological responses of TBI and promote recovery. To this end, numerous strategies, particularly pharmacological, have been investigated [7]. Although these functional consequences are important aspects of TBI morbidity, the current study focuses on the biological responses to TBI and post-injury treatment, rather than behavioral outcomes. However, despite significant benefits in preclinical studies, successful translation to the clinic has been inconsistent [8–10]. This limited translation underscores the need for novel therapeutic approaches as well as a better understanding of the biological factors that influence TBI pathology and treatment response. One such factor is biological sex, which is an important determinant of the response to TBI, yet relatively few studies have examined whether sex influences therapeutic efficacy [8–10]. Understanding this biological factor may be important for successful clinical translation, utilizing non-pharmacological therapeutic approaches.

Human neural stem cells (NSCs) are attractive candidates for restoring brain function after injury through both reconstruction (cell replacement) and repair (neuroprotection). Their inherent pathotropism toward sites of central nervous system (CNS) damage enables them to deliver therapeutic benefit by secreting growth factors, suppressing inflammation, reducing axonal injury, and differentiating into mature neural cell types. NSCs are currently under investigation in preclinical and clinical studies for stroke, multiple sclerosis, and other neurodegenerative conditions [11, 12]. In oncology, NSCs have been engineered to deliver a range of anti-cancer agents and have demonstrated therapeutic efficacy in models of primary and metastatic brain tumors [13, 14]. Despite these advances, a central challenge in NSC-based therapy remains the efficient delivery of sufficient, viable cells to achieve therapeutic effect within the CNS [13–15]. To address this challenge, multiple delivery routes of NSCs, including intravenous, intracranial, and intranasal (IN), have been evaluated [15, 16]. Preclinical studies further support the therapeutic potential of NSCs in reducing lesion volume and improving functional outcomes after TBI [17].

We developed human NSC lines engineered to stably express L-MYC (LMNSCs), which exhibit favorable growth properties, multi-lineage differentiation potential, and robust neuroprotective activity in vitro and in vivo [18, 19]. These LMNSCs and their extracellular vesicles (EVs) have been extensively characterized using scRNA-seq, RNA-seq, and proteomics, enabling identification of the most potent neuromodulatory line, LMNSC01 [16, 20, 21]. Here, we investigated sex-specific responses to controlled cortical impact (CCI) injury and subsequent LMNSC01 treatment. We found that a single IN administration of LMNSC01 was sufficient to modulate inflammatory and immune signaling molecules (*Tgfb1, Ccl9*, *Ccl21a* and *Xcr1*) and upregulate endogenous anti-inflammatory mechanisms (*Il1rn*) after TBI in male rats. In female rats, LMNSC01 treatment significantly decreased expression of genes related to macrophage activation, MHC presentation, and neuroinflammation induced by injury in female rats (*Il18*, *Pik3cg*, *Tgfbr2*, *Abca1*, *Irak4*, and *Cd74*). Cytokine ELISA revealed the LMNSC01 treatment modulated proteins associated with neuroinflammation and neurovascular remodeling (ICAM-1 and CX3CL1) in both male and female rats. Collectively, these findings demonstrate that a single IN dose of LMNSC01 reprograms the post-injury inflammatory microenvironment, promotes tissue regeneration, and supports neurovascular repair, highlighting its potential as a minimally invasive therapeutic strategy for TBI.

## Methods

### Animals

All rat studies were performed according to ARRIVE (Animal Research: Reporting In Vivo Experiments) guidelines published in 2010 (https://arriveguidelines.org). Adult male (n = 41) and female (n = 29) rats (Sprague–Dawley; Envigo RMS, Inc., Indianapolis, IN) were pair-housed in ventilated polycarbonate micro-isolator rat cages and maintained in a temperature-controlled (21 ± 1 °C) and light-controlled (0700–1900 h) vivarium with *ad libitum* food and water access. The rats were allowed to acclimate for 1 week before undergoing experimental procedures. After the acclimatization period, the rats were randomly assigned to brain injury groups (male CCI + LMNSC01 [n = 12], female CCI + LMNSC01 [n = 10], male CCI + vehicle [Veh, n = 11], and female CCI + Veh [n = 9]) and control groups (male Sham + LMNSC01 [n = 9], female Sham + LMNSC01 [n = 5], male Sham + Veh [n = 9], and female Sham +Veh [n = 5]) to assess experimental differences among the surgery groups ± LMNSC01 cells. All experimental manipulations were conducted during the lights-on phase. All procedures were conducted in accordance with the Guide for the Care and Use of Laboratory Animals and were reviewed and approved by the University of Pittsburgh Institutional Animal Care and Use Committee (IACUC). Mindful effort was made to minimize animal pain, suffering, or discomfort and to limit the number of rats used.

### Controlled Cortical Impact Surgery

Anesthesia was induced with 4% isoflurane in 2:1 N_2_O:O_2_, which was followed by endotracheal intubation, placement on a stereotaxic frame, and mechanical ventilation. Anesthesia was maintained with 2% isoflurane and the same volume of carrier gases. Core temperature was monitored and maintained at 37 ± 0.5 °C with a rectal thermistor and heating pad, respectively. Utilizing aseptic procedures, a CCI injury was produced as previously described [22, 23]. Specifically, a midline scalp incision was made, the skin and fascia were reflected, and a craniectomy was made above the right parietal-hemisphere with a high-speed dental drill. The bone flap was removed, and the craniotomy was enlarged using bone rongeurs to ensure accurate placement of the impact tip (6 mm, flat). Following the craniectomy, the impact tip was lowered until contact with the dura mater (zero point). The impact tip was then retracted into a ready position and set to impact 2.8 mm below the surface of the dura. Impact speed was set to 4 m/s with a dwell time of 100 ms. Following impact, anesthesia was discontinued to reduce neuroprotective confounds, and the rats were extubated, rapidly sutured, and assessed for acute neurological outcomes to ensure brain injury equivalence across all groups. Sham rats were not subjected to the impact but received all other surgical manipulations to control the potential effects of anesthesia and craniectomy.

### Cyclosporin A, LMNSC01 cells, and Veh administration

To facilitate the engraftment of LMNSC01 cells, an immunosuppressive regimen of cyclosporin A (CsA; 10 mg/kg) was provided subcutaneously to all rats beginning 2 days before the first IN administration of LMNSCs or Veh and continuing once per day until the brains were harvested. Human LMNSC01 cells from our Master Cell Bank were prepared for IN administration according to established SOPs [18, 19, 21, 24]. Briefly, LMNSC01 cells were thawed, washed, and re-suspended in 2% Human Serum Albumin (HSA) in PBS (6 × 10^6^ cells in 24 μL for a single bolus dose administration). Prior to infusion, rats were anesthetized using Isoflurane and placed on a nose cone in the supine position. The nasal epithelium was disrupted via hyaluronidase infusions at 6 μL per nostril at 2 min intervals for a total volume of 24 μL. After 20 min of disruption exposure, rats then received IN infusions of LMNSC01 cells or the same volume of Veh (6 μL per nostril drop at 2 min intervals for a total volume of 24 μL). Veh consisted of 2% HSA in PBS. IN dosing occurred on day 7 after CCI or Sham injury.

### Euthanasia and tissue collection

On postoperative day 39, rats were deeply anesthetized with Fatal-Plus® (0.3 mL, intraperitoneal, Henry Schein Animal Health; Warrendale PA). For NanoString and ELISA analysis, brain tissue was collected after euthanasia, rapidly dissected, and SNAP frozen in liquid nitrogen for storage at -80 °C.

### NanoString nCounter gene expression analysis

A subset of rats was randomly selected NanoString nCounter® gene expression analysis. Male cohorts included TBI and contralateral (CL) tissue from CCI + LMNSC01 (n = 5), CCI + Veh (n = 5), and Sham + LMNSC01 (n = 6), and Sham + Veh (n = 6). Female cohorts included TBI tissue from CCI + LMNSC01 and CCI + Veh groups (n = 7/group), CL tissues from the same injury groups (n = 5/group), and both TBI and CL tissues from Sham + LMNSC01 (n = 4) and Sham + Veh (n = 3) controls.

Total RNA from snap frozen brain tissue was profiled using the mouse nCounter® PanCancer Immune Profiling Panel for the NanoString nCounter Platform (CAT#Catalog#XT-CSO-MIP1-12, NanoString Technologies). The nCounter® immune profiling panel allows for the quantification of 770 genes related to immunological functions and housekeeping genes for normalization [25]. Purified RNA was hybridized with the panel-specific Reporter CodeSet and Capture CodeSet according to the manufacturer’s instructions. Excess probes were removed, and the reporter-capture complex was quantified using the nCounter® Digital Analyzer (NanoString Technologies, Seattle, WA). Raw RCC files were analyzed using the HyperScale architecture within ROSALIND® (Version 3.39.28.7), developed by ROSALIND, Inc. (San Diego, CA, USA). Normalization was conducted by dividing raw counts within each lane by the geometric mean of selected normalizer probes from the same lane. Housekeeping genes used for normalization were selected using the geNorm algorithm implemented in the NormqPCR R package. Differential expression statistics, including fold change and p-values, were calculated using the “fast method” described in the nCounter® Advanced Analysis 2.0 User Manual. Data normalization and differential gene expression analysis were performed using default parameters.

Principal component analysis (PCA) was performed in R (v4.5.2) using normalized count data exported from ROSALIND®. Pathway and functional enrichment analyses were performed using a hypergeometric distribution framework. The topGO R package was used to account for local similarities and dependencies among Gene Ontology (GO) terms using the Elim pruning method. Enrichment was calculated relative to an experiment-specific background gene set referencing Interpro [27], NCBI [28], MSigDB [29, 30], REACTOME [31], and WikiPathways [32] annotation databases. Canonical pathway and network analyses were further performed using QIAGEN Ingenuity Pathway Analysis (IPA; QIAGEN Inc., Redwood City, CA, USA). Volcano plots and bar charts of pathway enrichment scores were generated in R (v4.5.2).

### Multiplex sandwich-based ELISA assay

Multiplexed ELISA assays measuring the concentration of cytokines were performed on brain tissue from both CCI (CCI + LMNSC01 and CCI + Veh) and Sham surgery (Sham + LMNSC01 and Sham + Veh) groups. The Quantibody® Rat Cytokine Array 67 Kit (CAT#QAR-CAA-67-1, RayBiotech Inc), which consists of two non-overlapping arrays (QAR-CYT-3 and QAR-CYT-4, RayBiotech Inc.) that quantitatively measure the concentration of 67 rat cytokines. Tissues lysates were prepared in 1× lysis buffer (EL-lysis, RayBiotech) and Protease inhibitor (AA-PI, RayBiotech) using a hand-held tissue homogenizer (PRO-PK-01200PMGXL, Pro Scientific) and shipped to RayBiotech Services.

For all samples, proteins that measured at a concentration less than the limit of detection (LOD), the lowest reliable estimate by the instrument, were ignored. Additionally, protein concentrations less than the LOD were replaced with the LOD to facilitate ratio calculation. Finally, protein concentrations greater than the maximum measurable concentration (Max) were replaced by the Max for consistency. The relative change in cytokines is the ratio of mean protein expression in 1) CCI + Veh vs Sham and 2) CCI + NSC vs CCI + Veh samples.

### IHC staining and quantification

Brain tissue from each male rat treatment group (Sham + Veh [n = 3], Sham + LMNSC01 [n = 3], CCI + Veh [n = 6], and CCI + LMNSC01 [n = 7], totaling n = 19) were cut and prepared on paraffin slides prior to IHC and hematoxylin & eosin (H&E) staining. Brain tissue was cut coronally (n=200 per rat) from 3 mm to 5 mm posterior from the bregma, and every 10th slide was stained with H&E. Paraffin brain sections were stained with rat specific markers IBA1 (CAT#019-19741;1:400, Fujifilm-Wako), CD68 (CAT#ab125212; 1:100, abcam), and CD31 (CAT#ab281583; 1:100, abcam).

## Results

### Differential gene expression reveals LMNSC01 treatment modulates genes and pathways in male TBI rats

To evaluate the impact of CCI injury and LMNSC01 treatment on recovery after TBI, adult immunocompetent male rats received CCI surgery on day 0, followed by single dose IN administration of LMNSC01 cells (6 × 10^6^ cells) or Veh on day 7 post-injury. Animals were monitored for 39 days then euthanized for molecular and histological analyses (**Fig. 1A**). Brain tissue collection was performed according to the experimental schema (**Fig. 1B**). PCA was performed on normalized gene expression profiles of TBI (**Fig. 1C**) and CL (**Fig. S1A**) tissue samples. Separation of most CCI + LMNSC01 samples from CCI + Veh samples indicated differences in gene expression profiles in response to LMNSC01 treatment.

**Figure 1.**
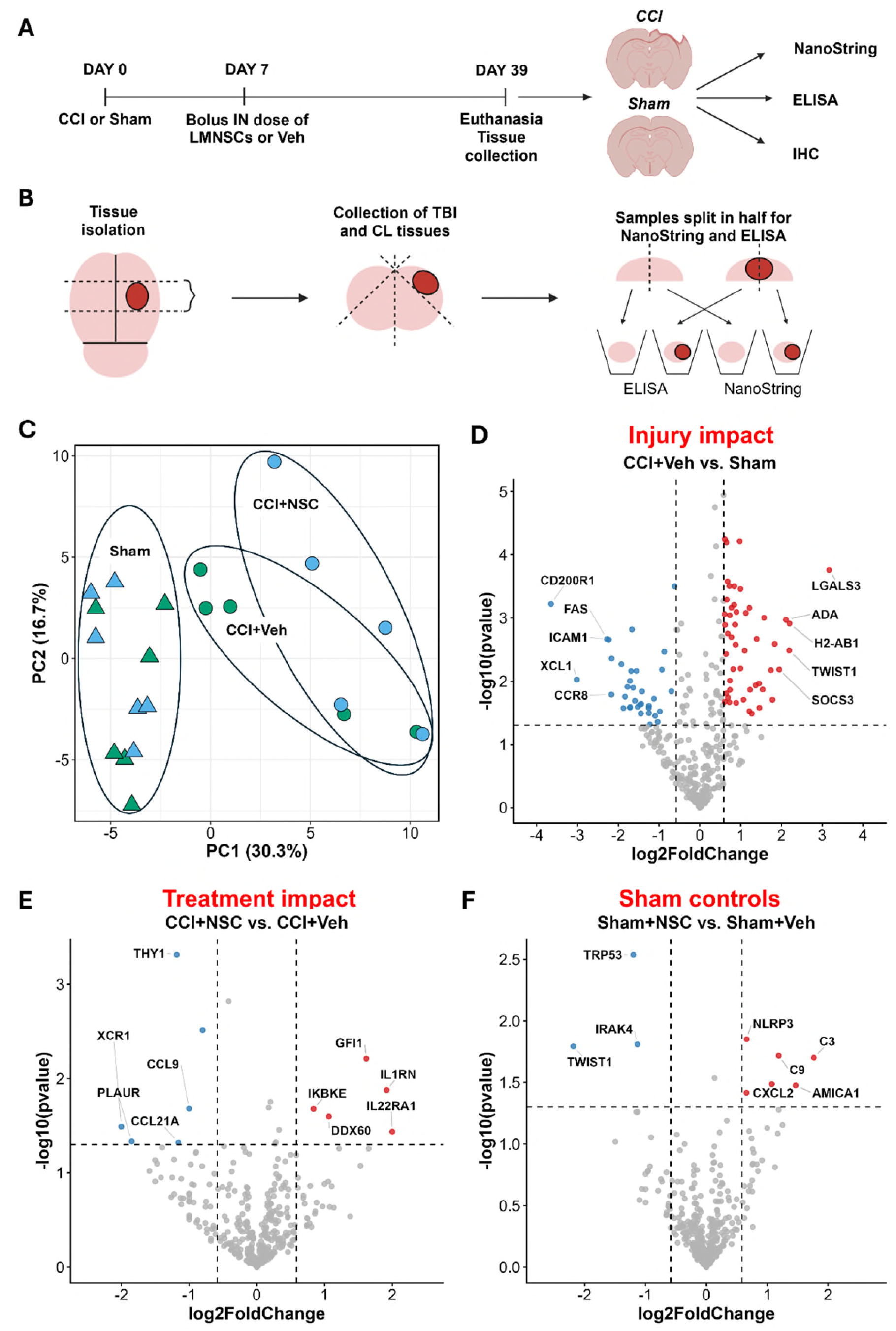
Gene expression analysis after CCI injury and CCI injury followed by IN LMNSC01 treatment in male rats. **(A)** Diagram of experimental timeline for IN administration of LMNSC01 cells following CCI injury in male rats. Seven days after CCI or Sham surgery, rats received a single IN injection of 6 × 10^6^ LMNSC cells or a Veh control. Brains were collected after 39 days for NanoString, ELISA, and IHC. **(B)** Schematic of tissue preparation for ELISA and NanoString analysis. **(D, F)** Differentially expressed genes (DEG) identified by NanoString analysis. **(C)** PCA clustering CCI + Veh, CCI + LMNSC01, Sham + Veh, and Sham + LMNSC01 samples based on gene expression profiles. **(D)** Volcano plot showing differences in TBI (right) side tissue gene expression for Veh-treated CCI surgery vs Sham surgery rats, with the x-axis showing log_2_ fold change and the y-axis showing the negative log_10_ p-value. Differential gene expression analysis was performed by ROSALIND. Red and blue circles indicate significantly upregulated (log₂FC ≥ 0.585, p < 0.05) and downregulated (log₂FC ≤ -0.585, p < 0.05) genes, respectively; top 10 genes are labeled. **(E, F)** Volcano plots as in (D), showing differences in TBI (right) side tissue gene expression for LMNSC01-vs Veh-treated CCI surgery rats.

To assess the impact of CCI injury on gene expression, differential gene expression analysis identified genes that were significantly up or downregulated (|log₂ Fold Change| ≥ 0.585, p < 0.05) in CCI + Veh vs. Sham groups (**Fig. 1D**). In TBI side tissue, the strongest increases in expression due to CCI injury were observed for *Lgals3* (LogFC = 3.17), *H2-Ab1* (LogFC = 2.20), *Twist1* (LogFC = 2.19), *Ada* (LogFC = 2.11), and *Socs3* (LogFC = 1.95). *Lgals3* and other significantly upregulated genes including *C3, C4b, Abca1,* and *Nlrp3*, are associated with inflammation, reflecting engagement of the innate immune response signaling after CCI Injury. Additional upregulated genes included *Pdgfc,* which is implicated in angiogenesis and increased blood–brain barrier (BBB) permeability, and *Ly96*, a TLR4 coreceptor involved in innate immune signaling. Increases in the *Stat1* and *Stat3* transcription factors reflect the activation of cytokine signaling cascades. In contrast, a marker of anti-inflammatory microglial subpopulations (*Cd200r*), cell death receptor (*Fas*), and a DNA damage response protein (*Cdkn1a*) were downregulated in CCI rats compared to Sham. These changes in gene expression are consistent with chronic neuroinflammation and cell cycle dysregulation after brain Gene expression in CCI rats treated with LMNSC01s was compared with Veh-treated controls to characterize the effect of IN LMNSC01 treatment after CCI (CCI + NSC vs CCI + Veh; **Fig. 1E**). *Il1rn*, an antagonist to the IL-1 receptor that inhibits Il1a and Il1b activities, was significantly upregulated following LMNSC01 treatment. Clinical trials testing recombinant Il1rn protein to treat TBI are currently in progress [26]. Additional upregulated genes, *Ddx60*, *Ikbke*, and *Gfi1*, are involved in Toll-like receptor and RIG-I– like receptor signaling and regulation of type I interferon pathways. *Gfi1* is responsible for regulating the inflammatory response to TLR signaling and represses inflammasome activity, suggesting a possible role for LMNSC01 treatment in controlling inflammation post-TBI [27]. LMNSC01-treated rats also exhibited increased expression of *Il22ra1*, which encodes the IL20, Il22, and Il24 cytokine receptor implicated in epithelial survival and tissue repair, suggesting activation of regenerative signaling mechanisms. Conversely, chemokine and chemokine receptor genes (*Ccl9* and *Xcr1*) involved in initiating inflammatory signaling cascades were downregulated following LMNSC01 treatment. Another chemokine, *Ccl21a*, which is expressed by injured neurons to activate microglia, was also downregulated following LMNSC01 treatment [28]. While *Tgfβ1* has neuroprotective effects in the acute phase of TBI, it can become a potent fibrotic factor when activated long term [29, 30]. LMNSC01-mediated deactivation of *Tgfβ1* may contribute to wound healing in the chronic phase of TBI. In contrast, differential gene expression analysis of Sham + LMNSC01 vs. Sham + Veh samples identified a modest upregulation of genes involved in innate immune signaling, including *Nlrp3, C3,* and C9, after LMNSC01 treatment in the absence of a TBI (**Fig. 1F**). Collectively, these data demonstrate that a LMNSC01 treatment enhances expression of immune regulatory genes while suppressing proinflammatory gene expression in the male rat brain after CCI.

To assess transcriptional responses distal to the injury site, differential gene expression CL brain tissue using the same comparison groups as for TBI tissue. Fewer transcriptional changes in response to CCI surgery were observed on the CL side compared to the TBI side (CCI + Veh vs. Sham; **Fig. S1A**). Significant decreases in expression of genes involved in T Cell functions (*Spp1, Tigit, Ctla1, Tnfsf14,* and *Eomes*) and innate immune signaling (*Cd35, Cxcr2, Derping1, Cd55,* and *Defb1*) after CCI injury on the CL side suggest that there may be an anti-inflammatory response that occurs away from the injury site after CCI injury. LMNSC01 treatment induced significant transcriptional changes in the CL hemisphere following TBI (CCI + LMNSC01 vs CCI + Veh; **Fig. S1C**), including downregulation of a natural killer (NK) cell and cytotoxic lymphocyte activation receptor (*Klra4*), along with adhesion molecules involved in leukocyte trafficking across the BBB (*Itgal* and *Vcam1*). Suppression of these genes in the CL hemisphere indicates that LMNSC01 treatment is associated with reduced neuroinflammatory gene expression beyond the primary CCI injury site in male rats.

### Differential gene expression reveals LMNSC01s modulate genes and pathways in female rats

Because biological sex can influence microglial activation, peripheral immune-cell recruitment, and inflammatory signaling following TBI, we next determined whether CCI and IN LMNSC01 treatment produced distinct transcriptional responses in female rats [31–33]. Transcriptional changes in immunocompetent female rats were assessed following CCI injury and IN LMNSC01 treatment using the NanoString immune gene expression platform used for the male cohort (**Fig 2A**). PCA of normalized gene expression profiles from the TBI site (**Fig. 2B**) shows clustering of Sham samples and some separation of CCI + NSC and CCI + Sham groups.

**Figure 2.**
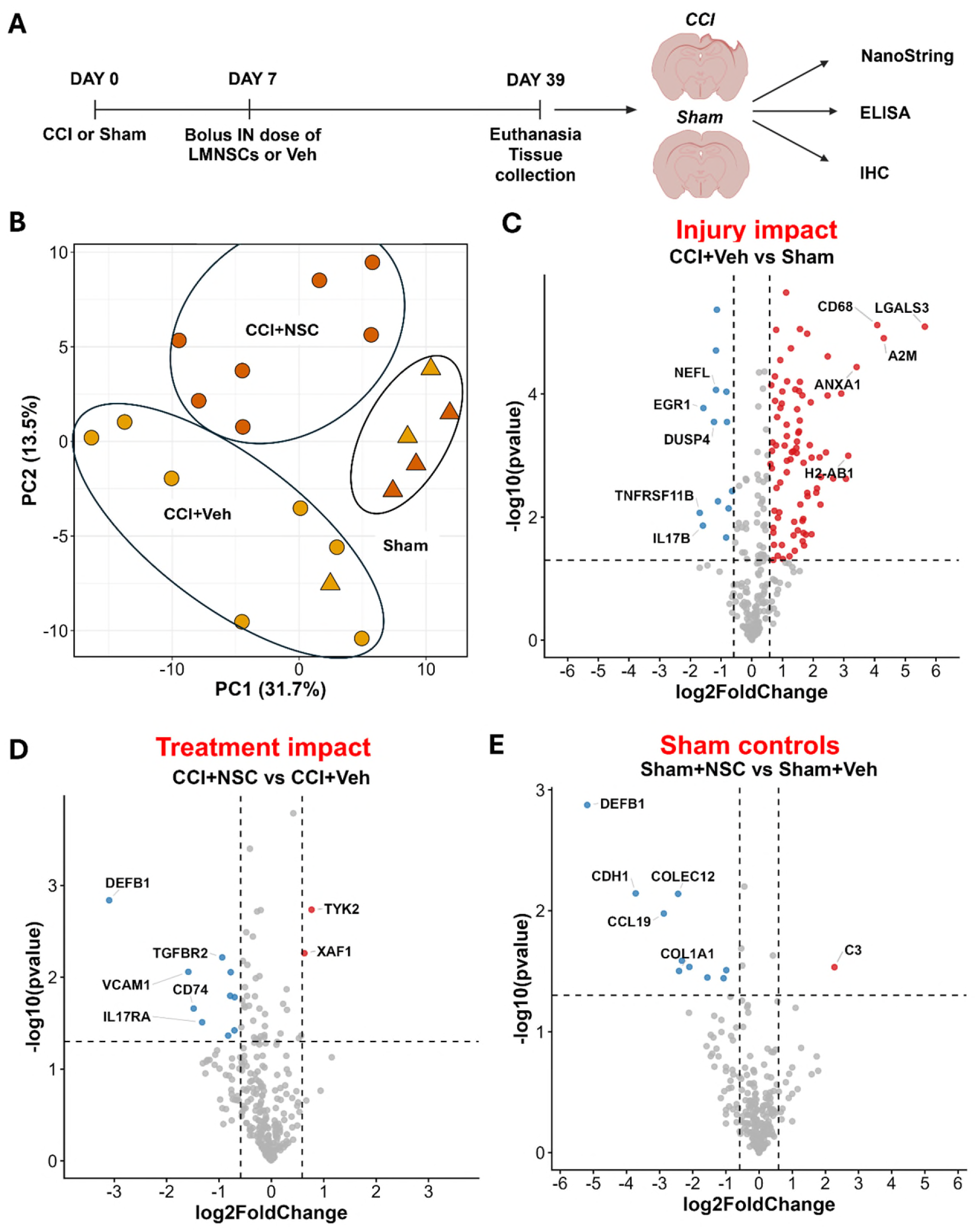
Gene expression analysis after CCI injury and CCI injury followed by IN LMNSC01 treatment in female rats. **(A)** Diagram of experimental timeline for IN administration of LMNSC01 cells following CCI injury in female rats. Seven days after CCI or Sham surgery, rats received a single IN injection of 6 × 10^6^ LMNSC cells or a Veh control. Brains were collected after 39 days for IHC, ELISA, and gene expression analysis. **(B, E)** DEG identified by NanoString analysis. **(B)** PCA clustering CCI + VEH, CCI + LMNSC01, Sham + Veh, and Sham + LMNSC01 samples based on gene expression profiles. **(C)** Volcano plot showing differences in TBI (right) side tissue gene expression for Veh-treated CCI surgery vs Sham surgery rats, with the x-axis showing log_2_ fold change and the y-axis showing the negative log_10_ p-value. Red and blue circles indicate significantly upregulated (log₂FC ≥ 0.585, p < 0.05) and downregulated (log₂FC ≤ -0.585, p < 0.05) genes, respectively; top 10 genes are labeled. **(D, E)** Volcano plots as in (C), showing differences in TBI (right) side tissue gene expression for LMNSC01-vs Veh-treated CCI surgery rats **(D)** and LMNSC01-vs Veh-treated Sham surgery rats **(E)**.

Differential gene expression analysis comparing CCI + Veh rats with Sham controls revealed a robust chronic neuroimmune response in female rats (**Fig. 2C**). Among the most strongly increased genes were *Lgals3* (log2FC = 5.64), *A2m* (Log2FC = 4.30), *Cd68* (log2FC = 4.09), *Anax1* (log2FC = 3.42), *H2-Ab1* (log2FC = 3.14), *Plau* (log2FC = 3.07), and *Cd74* (log2FC = 2.91). Additional increases in *Mrc1*, *Fcer1g*, *Slc11a1*, *Ctss*, and *Pik3cg* further supported a prominent microglial and leukocyte response at the chronic post-injury time point. Upregulation of *H2-Ab1*, *Cd73, H2-Eb1, Fcer1g,* and *Ctsh*, indicate enrichment of genes related to MHC presentation and increases in interferon induced proteins (*Ifi44, Ifitm, Ifih1,* and *Ifi35*) indicate increased interferon signaling after CCI in female rats. These findings suggest persistent activation and/or enrichment of myeloid and peripheral immune cell activity in the female brain following CCI.

We next determined the effect of LMNSC01 treatment after CCI injury in female rats by comparing immune response related gene expression in CCI + LMNSC01 with CCI + Veh groups (**Fig. 2D**). Several genes that were increased following CCI were subsequently reduced after LMNSC01 treatment, including *Il18*, *Pik3cg*, *Tgfbr2*, *Abca1*, *Irak4*, and *Cd74*, demonstrating reversal of specific components of the injury-associated transcriptional response. In addition to its involvement in macrophage activation, *Pik3cg* expression in neurons after TBI has been shown to induce endoplasmic reticulum stress leading to neuronal cell death. Importantly, *Pik3cg* knockout alleviated white matter injury and improved functional outcomes in a mouse CCI model [34]. LMNSC01-mediated reduction of *Il18* further suggested suppression of persistent inflammasome-associated cytokine signaling. In parallel, decreased *Cd74*, which encodes the invariant chain associated with MHC class II antigen presentation, was consistent with decreases in antigen-presentation that were strongly induced following CCI. Reduced *Il17ra* provided additional evidence of diminished inflammatory cytokine signaling. LMNSC01 also reduced *Vcam1*, which encodes an inducible endothelial adhesion molecule involved in leukocyte attachment and transmigration across activated vascular endothelium. This finding, together with a reduction in *Pik3cg*, suggests LMNSC01 treatment may deactivate neurovascular activation and leukocyte trafficking associated with TBI pathogenesis. Reduced *Tgfbr2*, receptor for *Tgf-β* ligands, may additionally reflect attenuation of neuroinflammation and extracellular matrix remodeling downstream of TGF-β. *Defb1* also decreased following LMNSC01 treatment, whereas *Tyk2* was increased; however, the functional implications of these changes in the context of TBI recovery remain to be determined.

Unlike in male rats, CCI injury produced very few gene expression changes on the CL side in female rats (**Fig. S2**). Upregulation of genes involved in inflammatory signaling, DNA damage response, and phagocytosis (*Il18, Prkcd,* and *Ncf4*) may reflect some injury related effects in the female rat brain away from the injury site (CCI + Veh vs Sham; **Fig. S2B**). Additionally, LMNSC01 treatment resulted in decreases in *Il18* and a T-cell activation gene, which may suggest possible anti-inflammatory effects distant from injured tissue (CCI + NSC vs. CCI + Veh; **Fig. S2c**).

### Pathway analysis reveals sex-dependent neuroimmune responses to CCI and LMNSC01 treatment in male and female rats

To elucidate sex-dependent pathways in gene expression changes for after CCI injury and LMNSC01 treatment, gene expression data of rats from all groups was subjected to functional analysis with NanoString immune response pathways. Broad increases in immune response categories were identified in male rats following CCI compared with Sham controls (CCI + Veh vs. Sham; **Fig. 3A**). Among the pathways with the greatest directed enrichment scores, CCI produced positive enrichment of pathways associated with Cell Cycle (DES = 1.66), Apoptosis (DES = 1.56), Adhesion (DES = 1.47), and Transporter Functions (DES = 1.17). Interestingly, Mast Cell Functions, Chemokines & Receptors, TNF Superfamily, and NK Cell Functions showed negative enrichment following CCI, indicating suppression of specific immune pathways rather than uniform activation at the chronic post injury time point. Decreases in Mast Cell Functions as well as Chemokines and Receptors pathways were also reflected in CL tissue after CCI (**Fig. S3A**). LMNSC01 treatment further modified pathways associated with CCI injury in male rats (CCI + NSC vs. CCI + Veh; **Fig. 3B**). The largest changes after treatment included negative directed enrichment of Dendritic Cell Functions (DES = -1.00), Senescence (DES = -0.88), and Chemokines & Receptors pathways (DES = -0.71).

**Figure 3.**
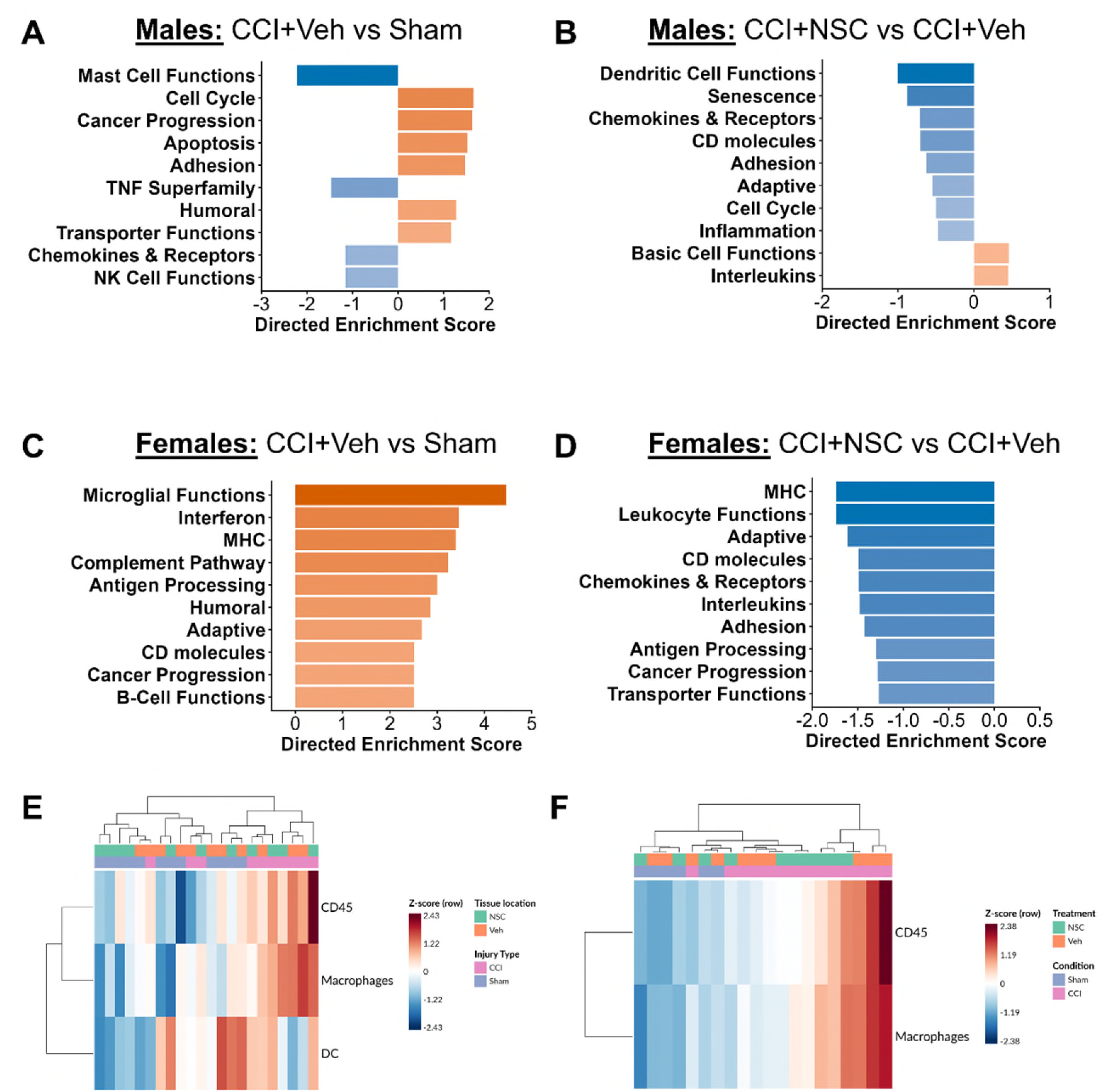
Sex differences in gene expression of brain tissue after CCI injury followed by intranasal LMNSC01 treatment in male and female rats. (A, **B)** Bar charts depicting NanoString functional enrichment pathway analysis of gene expression in TBI side (right hemisphere) brain tissue of male rats. The top ten significantly enriched pathways were identified using Gene Set Analysis in ROSALIND® (HyperScale architecture) with NanoString annotation databases. Treatment groups comparison: CCI + Veh vs Sham, CCI + NSC vs CCI + Veh. **(C, D)** Nanostring functional enrichment pathway analysis of gene expression in TBI side brain tissue of female rats. The top ten significantly enriched pathways were identified using Gene Set Analysis in ROSALIND®. Treatment groups comparison: CCI + Veh vs Sham, CCI + NSC vs CCI + Veh. **(E, F)** Cell type analysis for all groups in male and female rats at TBI sites.

Adhesion and Cell Cycle pathways that were activated after CCI injury show negative enrichment after LMNSC01 treatment. In contrast, Basic Cell Functions and Interleukins showed positive enrichment. Additional decreases in NK Cell Functions, Interferon signaling, and Complement Pathway were modulated after LMNSC01 treatment in CL tissue (Fig. S3B). These findings suggest that LMNSC01 treatment does not suppress immune signaling but selectively reduces pathways associated with cellular adhesion, immune cell surface molecules, and chemokine trafficking while maintaining or enhancing some cytokine signaling pathways. This pattern is consistent with the gene-level induction of endogenous immune regulatory molecules such as *Il1rn* following LMNSC01 treatment.

In female rats, functional pathway analysis revealed robust activation of neuroimmune pathways following CCI compared with Sham controls (Fig. 3C). Among the 10 pathways with the greatest absolute directed enrichment scores, Microglial Functions showed the strongest positive enrichment (DES = 4.46), followed by Interferon (DES = 3.46), and MHC (DES = 3.39). Positive enrichment was also observed for Complement Pathway (DES = 1.87), Antigen Processing (DES = 1.80), Humoral immunity (DES = 1.74), and Adaptive immunity (DES = 1.70), indicating further engagement of both innate and adaptive immune responses after CCI. Engagement of Adaptive immune signaling, Interferon, and MHC pathways was also reflected in CL tissue after CCI (**Fig. S3C**). LMNSC01 treatment produced a negative shift in many immune pathways associated with injury in female rats (CCI + LMNSC01 vs. CCI + Veh; **Fig. 3D**). Among the pathways with the greatest absolute directed enrichment scores, MHC showed the strongest negative enrichment (DES = −1.74), followed by Leukocyte Functions (DES = −1.74), and Adaptive immunity (DES = −1.61). Importantly, several pathways strongly increased following CCI (MHC, Antigen Processing, Adaptive immunity and CD Molecules), shifted toward negative enrichment following LMNSC01 treatment. Negative enrichment of Adaptive immunity, CD molecules, Interleukins and Inflammation in CL tissue after LMNSC01 treatment may reflect anti-inflammatory LMNSC01 mediated changes distal from the injury site.

Cell Type Profiler analysis (ROSALIND®) of TBI tissue identified distinct dendritic cell (DC), CD45-positive leukocyte, and macrophage-associated transcriptional signatures across all groups in male rats (**Fig. 3E**) and female rats (**Fig. 3F**). Hierarchical clustering demonstrated substantial inter-animal heterogeneity in both sexes; however, a subset of female CCI + Veh animals showed strong enrichment of CD45-positive leukocyte and macrophage signatures compared to CCI + NSC and Sham + Veh controls (**Fig. 3E**). Male samples generally exhibited similar levels of CD45-positive leukocyte and macrophage enrichment in CCI + Veh and CCI + NSC animals (**Fig. 3F**). The median log_2_ abundance scores for macrophages and CD45+ leukocytes but not dendritic cells were higher in both CCI + Veh and CCI + LMNSC01 groups of male (**Fig. S4A, B**) and female (**Fig. S4C-E**) rats. Dendritic cell signature mRNA was not detected in female rats. This is reflected in representative multiplex immunohistochemistry images, where staining for CD68 and IBA1 reveal increased macrophage/microglia presence in both CCI + Veh and CCI + LMNSC01 brain sections compared to Sham controls (**Fig. S5**).

### Canonical pathways after CCI injury and LMNSC01 treatment in male and female rats

Additional functional analysis using IPA compared activation Z-scores of canonical pathways in CCI + Veh vs. Sham and CCI + NSC vs. CCI + Veh in male and female rats. The top 40 most activated pathways when sorted by Z-score reveal broad activation of inflammatory and immune signaling pathways such as Interlekin-1 family signaling, T-cell receptor signaling, and NF-kB signaling following CCI in both sexes (**Fig. 4**). In parallel, pathways associated with tissue remodeling and neurovascular regulation, including PDGF, ILK, Apelin endothelial, and Rho GTPase signaling, were also enriched after injury. LMNSC01 treatment largely resulted in negative enrichment of these CCI-induced pathways. Both sexes exhibited a highly similar overall response to CCI and LMNSC01 treatment; many of the same major inflammatory pathways were activated by injury and subsequently deactivated by LMNSC01. However, the magnitude of activation appears greater in females for several immune-related pathways, including IL-1 signaling, NF-κB signaling, and adaptive immune pathways, indicating a somewhat stronger neuroimmune response to injury in female rats compared to male rats. Males and females both demonstrate reversal of these pathways following LMNSC01 treatment, suggesting that the therapeutic mechanism is largely conserved across sexes.

**Figure 4.**
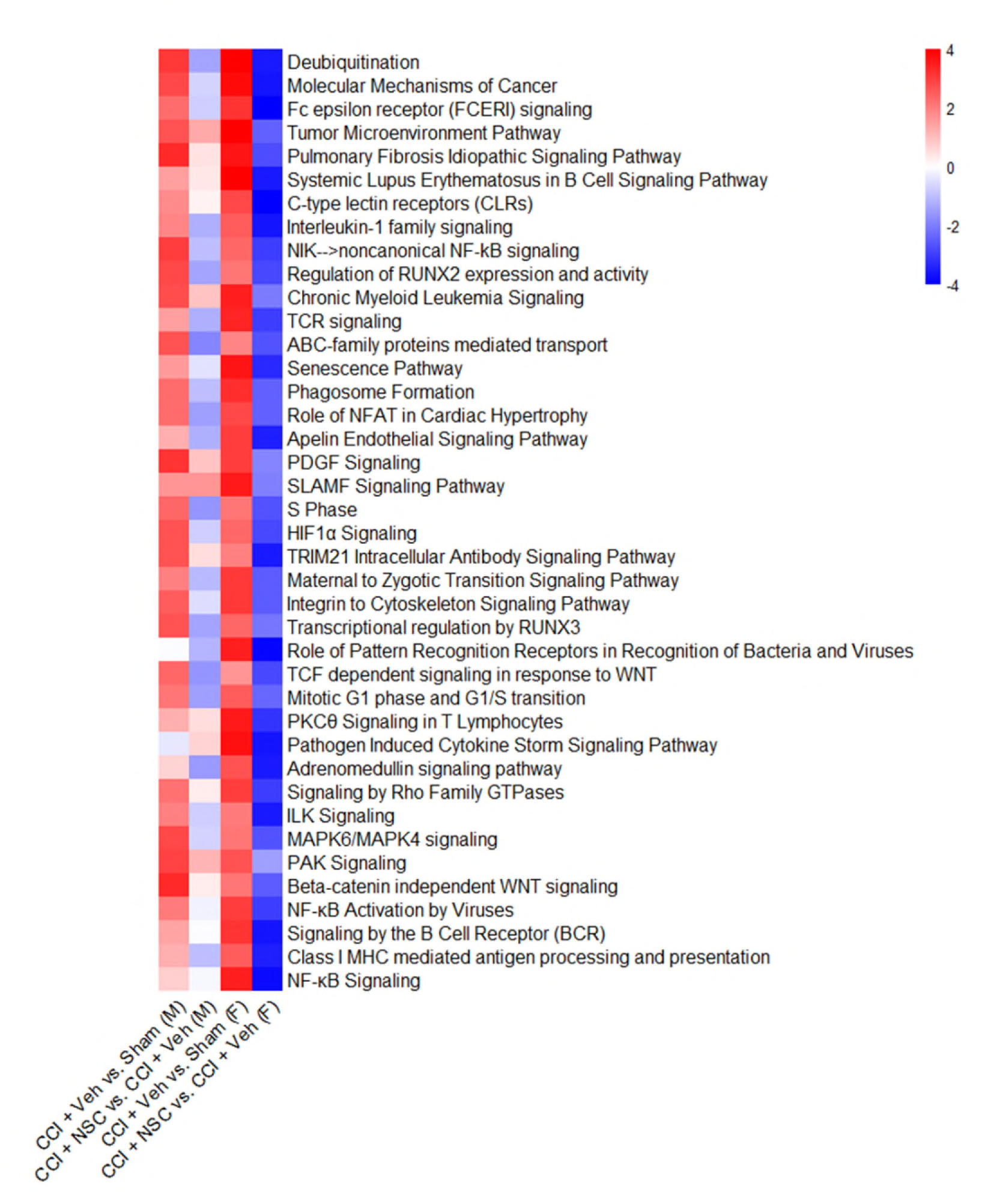
Affected canonical pathways after CCI injury and LMNSC01 treatment in male and female rats. Heatmap depicting canonical pathways identified by Ingenuity Pathway Analysis in CCI + Veh vs Sham and CCI + NSC vs CCI + Veh comparisons for both male and female rats. Top 40 pathways sorted by Z Score are displayed; red signifies activation, blue signifies repression.

### LMNSC01-mediated restoration of neurovascular and neuroimmune signaling through ICAM-1 and CX3CL1 after TBI in male and female rats

To determine the effects of CCI and LMNSC01 treatment on neuroinflammatory and neurovascular signaling, protein expression was quantified by multiplex tissue ELISA in snap-frozen brain tissue from male and female rats. Protein expression profiles were compared between CCI + Veh and all Sham surgery groups to define injury associated changes separately in male and female rats (**Fig. 5A, B**) and between CCI + NSC and CCI + Veh animal groups to determine treatment associated changes in each sex (**Fig. 5C, D**).

**Figure 5.**
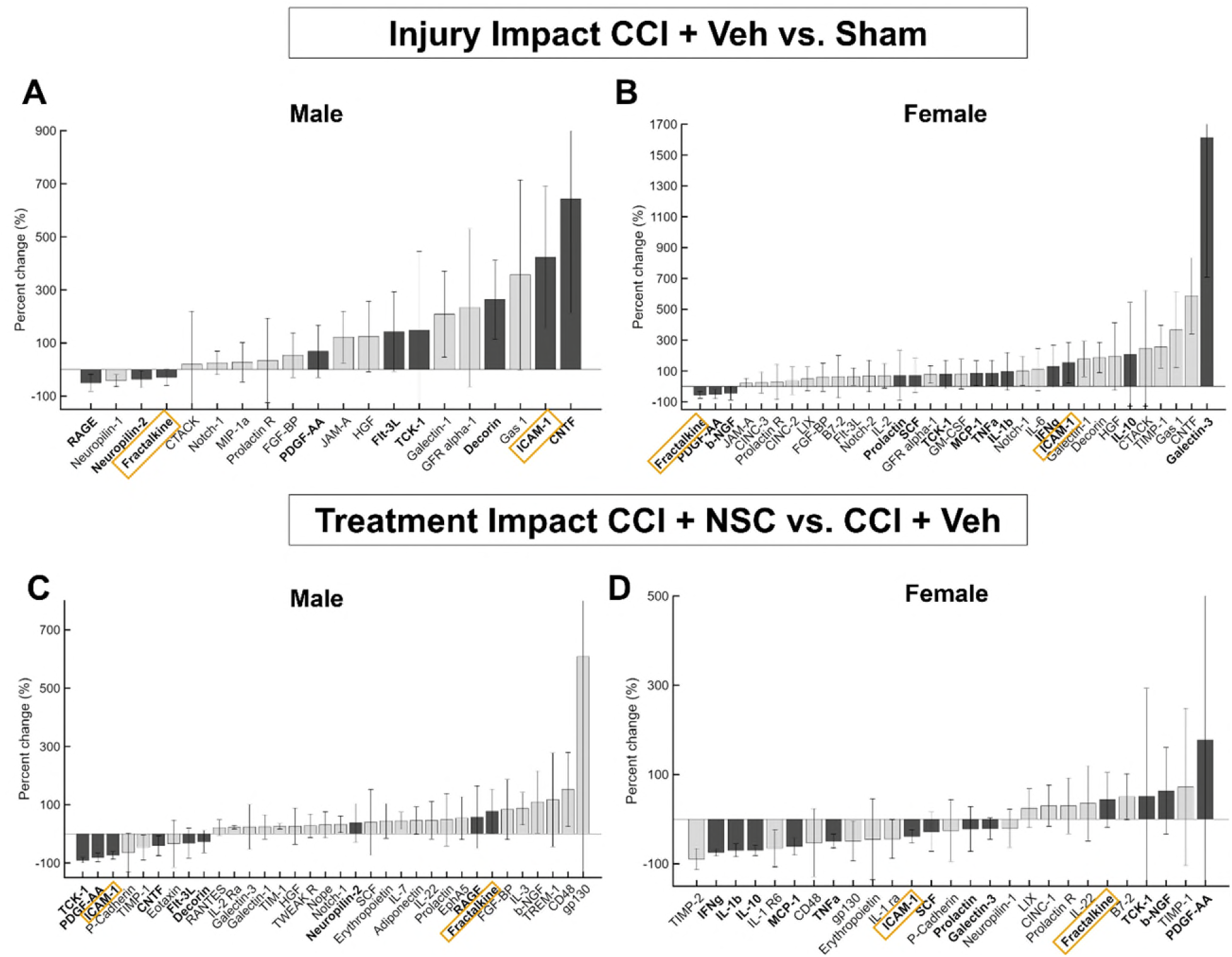
Sex differences of brain tissue protein expression after CCI and LMNSC01 treatment. **(A, B)** Bar charts showing percent change in protein concentration in male and female rat brain lysates of TBI side tissue from CCI + Veh vs. Sham groups measured by multiplex ELISA. Proteins with a percent change greater than ±20% are displayed. **(C, D)** Bar charts showing percent change in protein concentration in TBI tissue from CCI + NSC vs. CCI + Veh groups. Highlighted proteins are those that are regulated (up-down-regulated) by injury **(A and B)** and after LMNSC01 treatment **(B and D)**.

Increased levels of endothelial activation, inflammatory signaling, and extracellular matrix remodeling proteins, including ICAM-1, TCK-1, PDGF-AA, JAM-A, Gas-1, Decorin, Galectin-1, and CNTF, were observed following CCI in male rats (**Fig. 5A**). In contrast, Fractalkine (CX3CL1), Neuropilin-1, and Neuropilin-2 were reduced after CCI (**Fig. 5A**). LMNSC01 treatment substantially modified changes in protein expression induced by CCI (CCI + NSC vs. CCI + Veh; **Fig. 5C**). Among the most notable treatment associated changes were reductions in PDGF-AA, TCK-1, and ICAM-1, consistent with diminished vascular and inflammatory signaling associated with CCI injury in both TBI (**Fig. 5CB**) and CL (**Fig. S6**). LMNSC01 treatment was also associated with increases in several proteins involved in neurotrophic and neuronal signaling, as well as tissue regeneration, including β-NGF, EphA5, Neuropilin-2, and gp130.

Similar changes in protein expression following CCI injury were observed in female rats (CCI + Veh vs. Sham; **Fig. 5B**). Galectin 3, which has been identified as orchestrating inflammatory responses after TBI, demonstrated the greatest increase in abundance after CCI injury [35]. The *Lgals3* gene, which codes for Galectin 3, also had the greatest increase in mRNA expression after CCI in female rats (**Fig. 2B**). Additionally, CCI increased the abundance of proteins involved in ECM signaling and regulation of cell growth (Gas 1, TIMP-1 and Decorin) as well as cytokines that exhibit pro- and anti-inflammatory effects after TBI (IL-6, IL-1b, IFNg and IL-10). LMNSC01 treatment resulted in decreased abundance of injury associated cytokines IL-1b, TNFa, and IFNg in both TBI (**Fig. 5D**) and CL tissue **(Fig. S6**). In both male and female rats, LMNSC01 treatment decreased levels of ICAM-1 and increased levels of CX3CL1 (**Fig. 5C, D**). ICAM-1 is important in leukocyte adhesion and transmigration during brain injury, and studies have shown ICAM-1 depletion to have neuroprotective effects in a male mouse TBI model [36]. CX3CL1 is essential to neuron-microglia communication and has established roles in regulating microglial activation. IN administration of recombinant CX3CL1 inhibited neuroinflammation and promoted M2 microglial polarization in a male mouse CCI model [37]. These findings suggest that LMNSC01 cells may reduce leukocyte recruitment to and inflammatory microglial activity in tissue affected by a TBI.

### Gene interaction networks reveal sex-dependent organization of the CCI response

To characterize gene regulatory networks that orchestrate the immune response to CCI, IPA was used to determine biological relationships between differentially expressed genes after CCI (CCI + Veh vs. Sham) in both male and female rats (**Fig. 6**). While CCI induced highly interconnected inflammatory and immune regulatory gene networks in both male and female animals, females generally exhibited greater activation of immune-related networks following injury (**Fig. 6**). Transcription factors *Jun*, *Stat1*, and *Stat3*, were the most highly interconnected nodes in both male and female rats, suggesting that shared upstream regulators may activate inflammatory signaling cascades in both sexes (**Fig. 6**). Top co-regulated genes in male rats were involved in macrophage activation (for example, *Abca1*, *Fcer1g*, and *Mrc1*), as well as IL12 signaling (*Clu*, *Col1a1*, *Irf1* and *Nfkb2*) and cytokine storm pathways (*Pdgfc, Stat1, Stat3,* and *Tgfb1*) (**Fig. 6A**). In female rats, top co-regulated genes were also involved in IL12 signaling and cytokine storm pathways in addition to glucocorticoid receptor signaling (*Myc*, *Nfat1*, and *Mapk11*) and fibrosis (*Col1a1, Col3a1*, and *Vim*) (**Fig. 6B**).

**Figure 6.**
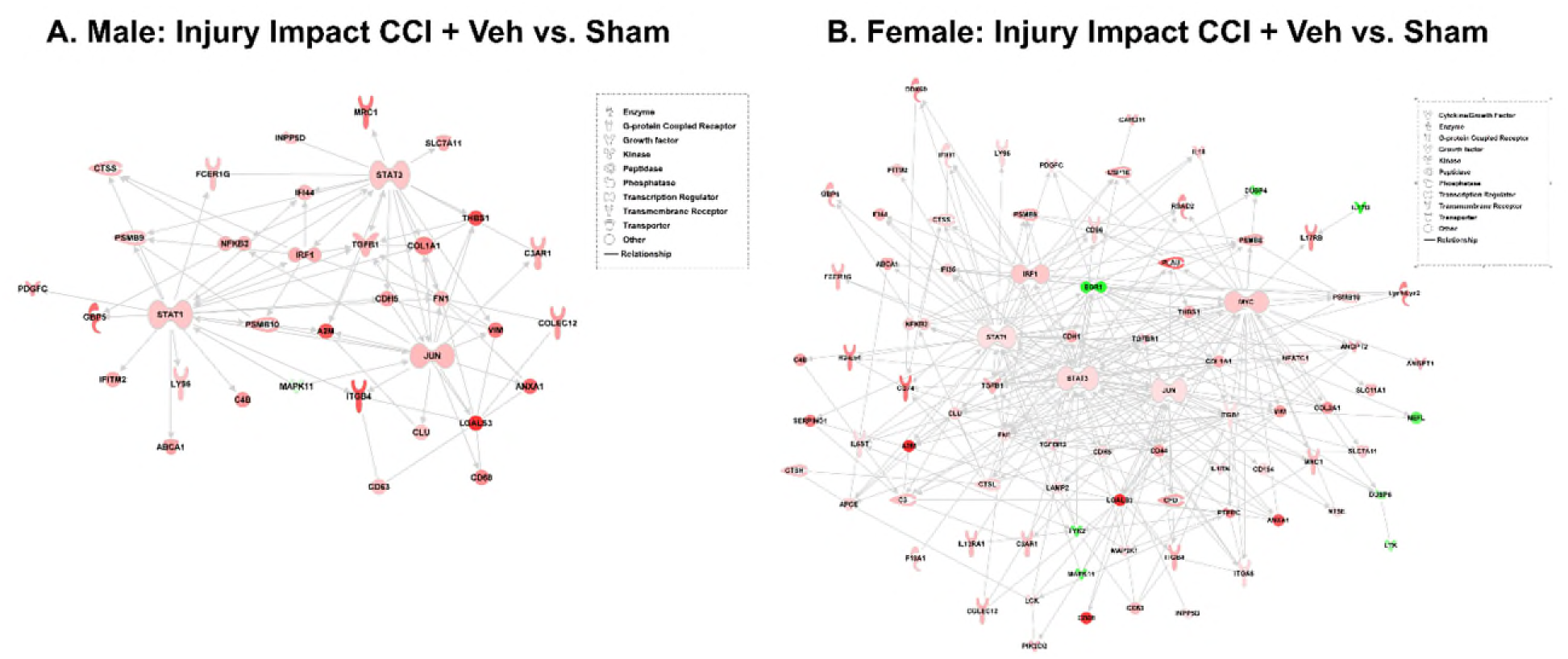
Interaction networks of brain injury-associated genes. **(A)** Network representation of significantly differentially expressed genes (|log₂FC | > 0.585, p < 0.05) in male rat TBI tissue samples from CCI + Veh vs Sham groups. Upregulated genes are shown in red; downregulated genes are shown in green. Lines indicate gene–gene interactions curated from the Ingenuity Pathway Knowledge base. **(B)** Network representation of injury-associated genes in female rat TBI tissue samples from CCI + Veh vs Sham groups as in (A).

## Discussion

TBI initiates a complex secondary injury response involving persistent neuroinflammation, neurovascular dysfunction, extracellular matrix remodeling, and disruption of homeostatic communication among neurons, glia, vascular cells, and infiltrating immune cells [38–40]. We hypothesized that NSCs are an attractive candidate to address TBI sequelae through their intrinsic ability to migrate to sites of brain inflammation, where they may support cellular regeneration by delivering growth factors, suppressing inflammation and differentiating into mature neuronal-lineage cells. Here, we demonstrate that a single IN administration of LMNSC01 cells 7 days after CCI produces sustained molecular changes in the injured brain, and that these effects occur in both male and female rats despite marked sex-dependent differences in the underlying response to injury.

In this study, we identified injury related increases in ICAM-1 protein abundance and subsequent decreases in abundance after LMNSC01 treatment following CCI in both male and female rats. ICAM-1 is an intercellular adhesion molecule involved in leukocyte and neutrophil transmigration across the endothelium and is thus an initiator of neuroinflammatory responses after TBI. ICAM-1 expression is induced by inflammatory cytokines upstream of NF-kB signaling cascades such as TNFa and IL-1b [41]. In this study, we found decreased levels of TNF and IL-1b and negative enrichment of NF-kB signaling pathways after LMNSC01 treatment of female rats after TBI, suggesting a possible mechanism through which LMNSC01 treatment may increase vascular integrity. Therapies targeting ICAM-1 are being investigated for TBI [36, 42].

In addition to persistent migration of peripheral immune cells across the BBB, chronic microglial activation has been implicated in neuroinflammation beyond the acute phase of TBI. The cytokine CX3CL1 maintains neuron-microglia homeostasis by limiting microglial activation and subsequent production of pro-inflammatory cytokines IL-1, IL-6, and TNFa through the CX3CL1-CX3CR1 axis [43, 44]. We found that CX3CL1, the abundance of which decreased after CCI injury, was increased by LMNSC01 treatment in CCI rats. Conserved increases in both male and female rats treated with LMNSC01 cells after TBI could suggest that NSCs may help reduce microglial polarization and neuroinflammation after TBI.

Sexual dimorphism in response to TBI has been reported in preclinical and clinical studies and may affect injury progression and treatment efficacy. In a review of clinical TBI studies, Gupte et al. found that the largest proportion of studies report women having worse outcomes [8]. The opposite was true for preclinical animal studies, with the largest proportion of studies reporting females having better outcomes. Interestingly, the percentage of studies reporting better outcomes in women and females increases with injury severity [8]. However, other clinical studies have found no gender-based difference in mortality or outcome favorability in patients with acute TBI [45]. Thus, it remains to be determined whether sex differences exist in TBI and what molecular mechanisms may account for them.

Pre-clinical studies have identified sex-dependent differences in microglial activation and infiltrating myeloid-cell responses following experimental TBI. Doran et al. identified greater infiltration of peripheral myeloid cells and proliferation of microglia at the injury site in adult male rats after CCI injury using flow cytometry [46]. However, our data indicates that female rats demonstrate increases in genes related to microglial activation and leukocyte infiltration after CCI injury, while cell cycle and apoptosis genes were more strongly enriched in male rats. Additionally, Scott et al. reported greater BBB permeability in female rats after CCI injury but no difference in microglial phenotype or inflammatory cytokine expression [9]. We identified increases in proteins and cytokines Galectin-3, TNF-a, and IL-1b after CCI in female rats that were not reflected in male rats, which could indicate sex-based differences in cytokine signaling after TBI. Responses to LMNSC01 treatment after CCI also varied between sexes; genes associated with leukocyte recruitment, antigen presentation, and inflammation (*Il18*, *Pik3cg*, *Tgfbr2*, *Abca1*, *Irak4*, and *Cd74*) that were upregulated after injury were significantly downregulated after LMNSC01 treatment in female rats, while LMNSC01 treatment resulted in the upregulation of endogenous immune regulators (*Il1rn*) and downregulation of chemokines (*Ccl9* and *Xcr1*) and *Tgfb1*, a major upstream regulator of inflammatory signaling after TBI. Together with our previous demonstration that intranasally administered LMNSC01 cells migrate toward sites of brain injury, these findings suggest that injury-targeted NSCs can engage distinct sex-dependent inflammatory environments yet converge on a shared neurovascular–neuroimmune reparative response. This extends the therapeutic paradigm for NSCs beyond cell replacement toward active modulation of the injured tissue microenvironment and provides support for further development of IN LMNSC01 as a minimally invasive regenerative therapy for TBI.

Several limitations should be considered when interpreting these findings. Although integrated transcriptomic, pathway, cell-type, and protein analyses identified coordinated LMNSC01-associated changes, these data primarily established molecular associations and did not demonstrate direct causality. In particular, the conserved reduction of ICAM-1 and restoration of CX3CL1 following treatment identified a candidate neurovascular–neuroimmune regulatory axis, but targeted manipulation of ICAM-1 and CX3CL1/CX3CR1 signaling will be required to establish their functional contribution to LMNSC01-mediated repair. Nevertheless, our findings demonstrate that a single, noninvasive IN administration of LMNSC01 reprograms persistent post-traumatic neuroimmune and neurovascular responses in both male and female rats. Despite distinct sex-dependent molecular responses to CCI and LMNSC01 treatment, both sexes converged on reduced ICAM-1 and restored CX3CL1, consistent with attenuation of endothelial activation and restoration of homeostatic neuron–microglia communication. Together, these findings identify restoration of neurovascular– neuroimmune homeostasis as a potential mechanism of LMNSC01-mediated repair and provide a strong rationale for further development of IN LMNSC01 as a minimally invasive regenerative therapy for TBI.

## Acknowledgments

This work was supported by National Institutes of Health grants P30CA033572, R01 Kline/Gutova 1R01NS121037-01, HD069620, NS099674, NS084967, NS113810 (AEK), NS094950, NS099683, NS110609 (COB), the Albert & Bettie Sacchi Foundation (MG), and the UMPC Rehabilitation Institute (COB). Research reported in this publication included work performed in the Research Pathology Services Shared Resources supported by the National Cancer Institute of the NIH under grant number P30CA033572. The content is solely the responsibility of the authors and does not necessarily represent the official views of the National Institutes of Health. The authors acknowledge the professional editing of Erin Keebaugh, PhD.

## Data Availability Statement

The NanoString nCounter gene expression data have been deposited in the Gene Expression Omnibus (GEO) under accession number **GSE322819**.

## Authors contributions

Conceptualization, AEK, MG, CB, and RR; methodology, BE, JPC, VA; validation, BE, IV; formal analysis, BE, IV, VA; investigation, BE, IV, VA; resources, AEK and MG; data curation, BE and RR; writing—original draft preparation, BE and MG; writing—review and editing, AEK, RR, CB, MG; visualization, JPC, IV, BE, RR; supervision, AEK and MG; project administration, AEK and MG; funding acquisition, RR, CB, AEK and MG. All authors have read and agreed to the published version of the manuscript.

