## Supplementary Figures for "Intranasal L-MYC Neural Stem Cells Restore Neurovascular and Neuroimmune Homeostasis After TBI via CX3CL1 Upregulation and ICAM-1 Downregulation"

**Corresponding Authors:**

**Margarita Gutova, PhD**

Assistant Professor

Beckman Research Institute at City of Hope

Department of Stem Cell Biology and Regenerative Medicine

1500 E. Duarte Rd.

Duarte, CA, 910101

Office: Miller Bld, Room 111

Lab: Kaplan Black Bldg, Room 109B

**Anthony E. Kline, PhD**

Professor

Physical Medicine & Rehabilitation,

Critical Care Medicine, Psychology, and Center for Neuroscience

Associate Director of Rehabilitation Research,

Safar Center for Resuscitation Research

University of Pittsburgh School of Medicine
 Children’s Hospital of Pittsburgh

John G. Rangos Research Center – Room 6126

Keywords: LMNSC neural stem cells, traumatic brain injury, controlled cortical impact brain injury, neurovascular, neuroimmune, CX3CL1, ICAM-1

**Supplementary Figures**

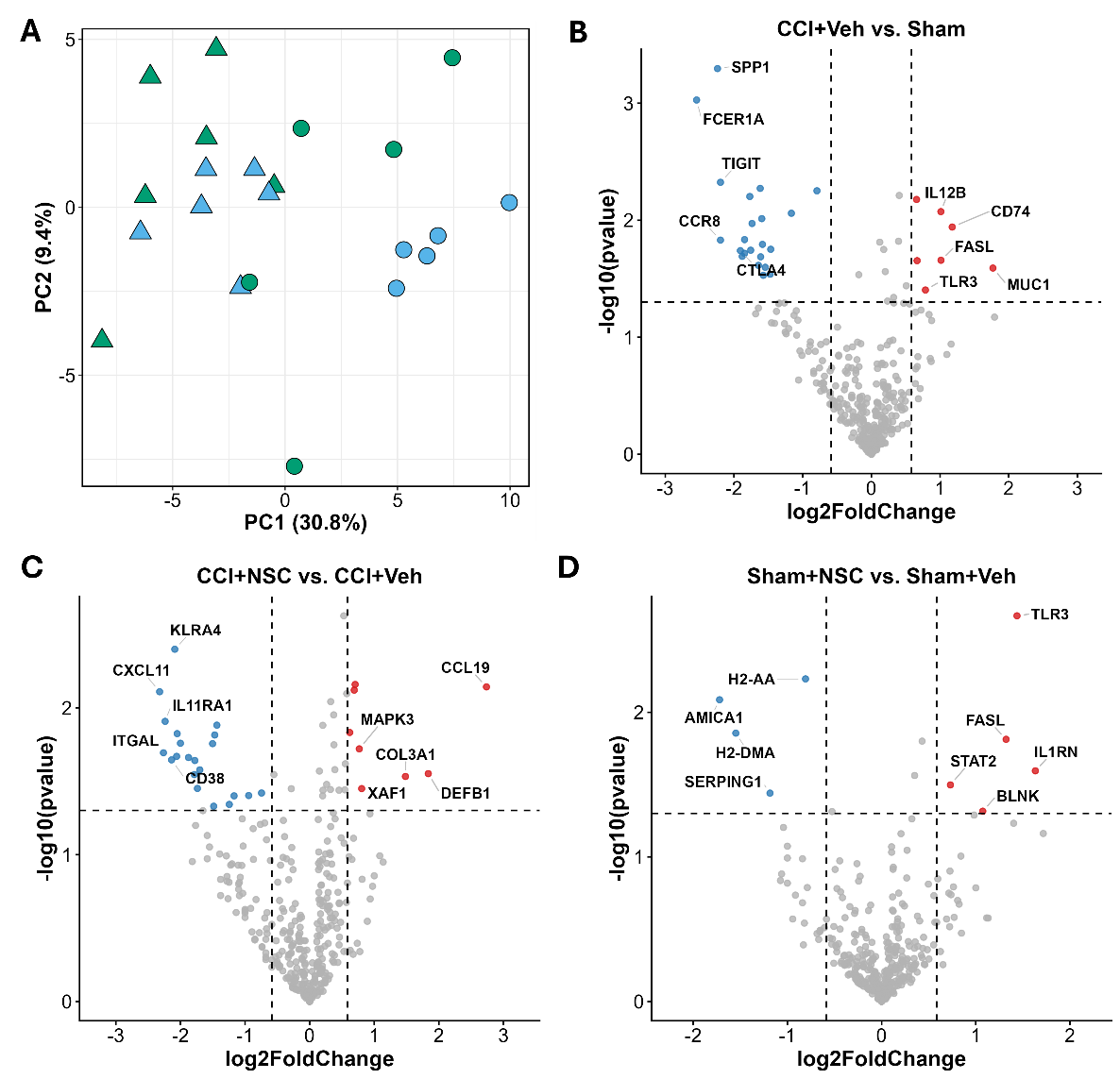

**Figure S1. NanoString analysis of contralateral side gene expression after CCI surgery and LMNSC01 treatment in male rats**

(A) Principal component analysis (PCA) clustering CCI (circles) and Sham surgery samples (triangles) treated with either LMNSC01s (blue) or a vehicle control (green).

(B) Volcano plot showing differences in CL (left) side tissue gene expression for vehicle treated CCI surgery vs sham surgery rats, with the x-axis showing log2 fold change and the y-axis showing the negative log10 p-value. Differential expression analysis was performed by ROSALIND. Red and blue circles indicate significantly upregulated (log₂FC ≥ 0.585, *p* < 0.05) and downregulated (log₂FC ≤ -0.585, *p* < 0.05) genes, respectively; top 10 genes are labeled.

(C, D) Volcano plots as in (B) showing differences in CL side tissue gene expression for LMNSC01 vs Vehicle treated CCI surgery rats (C) and LMNSC01 vs Vehicle treated Sham surgery rats (D).

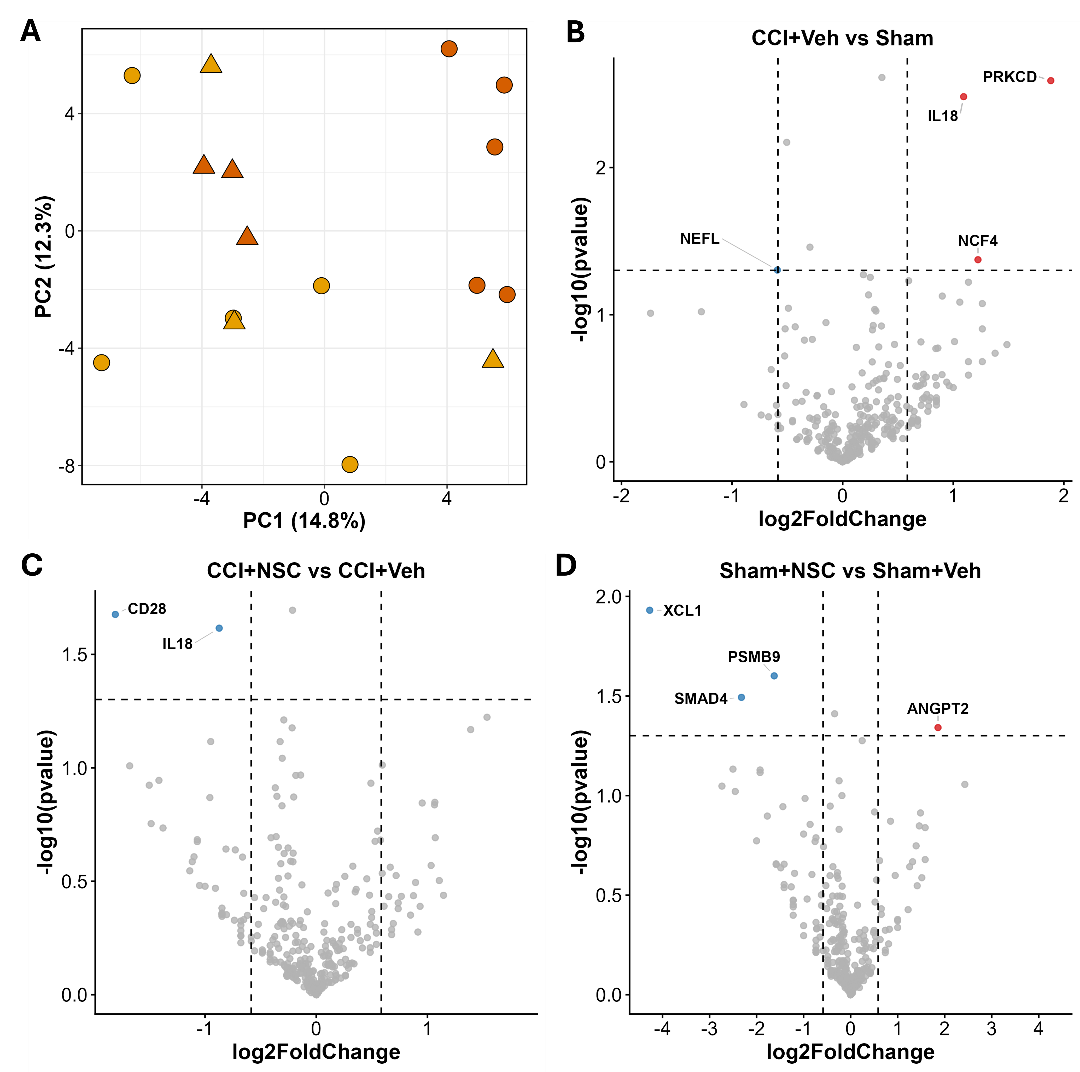

**Figure S2. NanoString analysis of contralateral side gene expression after CCI surgery and LMNSC01 treatment in female rats**

(A) Principal component analysis (PCA) clustering CCI (circles) and SHAM surgery samples (triangles) treated with either LMNSC01s (orange) or vehicle control (yellow).

(B) Volcano plot showing differences in CL (left) side tissue gene expression for vehicle treated CCI surgery vs sham surgery rats, with the x-axis showing log2 fold change and the y-axis showing the negative log10 p-value. Differential expression analysis was performed by ROSALIND. Red and blue circles indicate significantly upregulated (log₂FC ≥ 0.585, *p* < 0.05) and downregulated (log₂FC ≤ -0.585, *p* < 0.05) genes, respectively; top 10 genes are labeled.

(C, D) Volcano plots as in (B) showing differences in CL side tissue gene expression for LMNSC01 vs Vehicle treated CCI surgery rats (C) and LMNSC01 vs Vehicle treated Sham surgery rats (D).

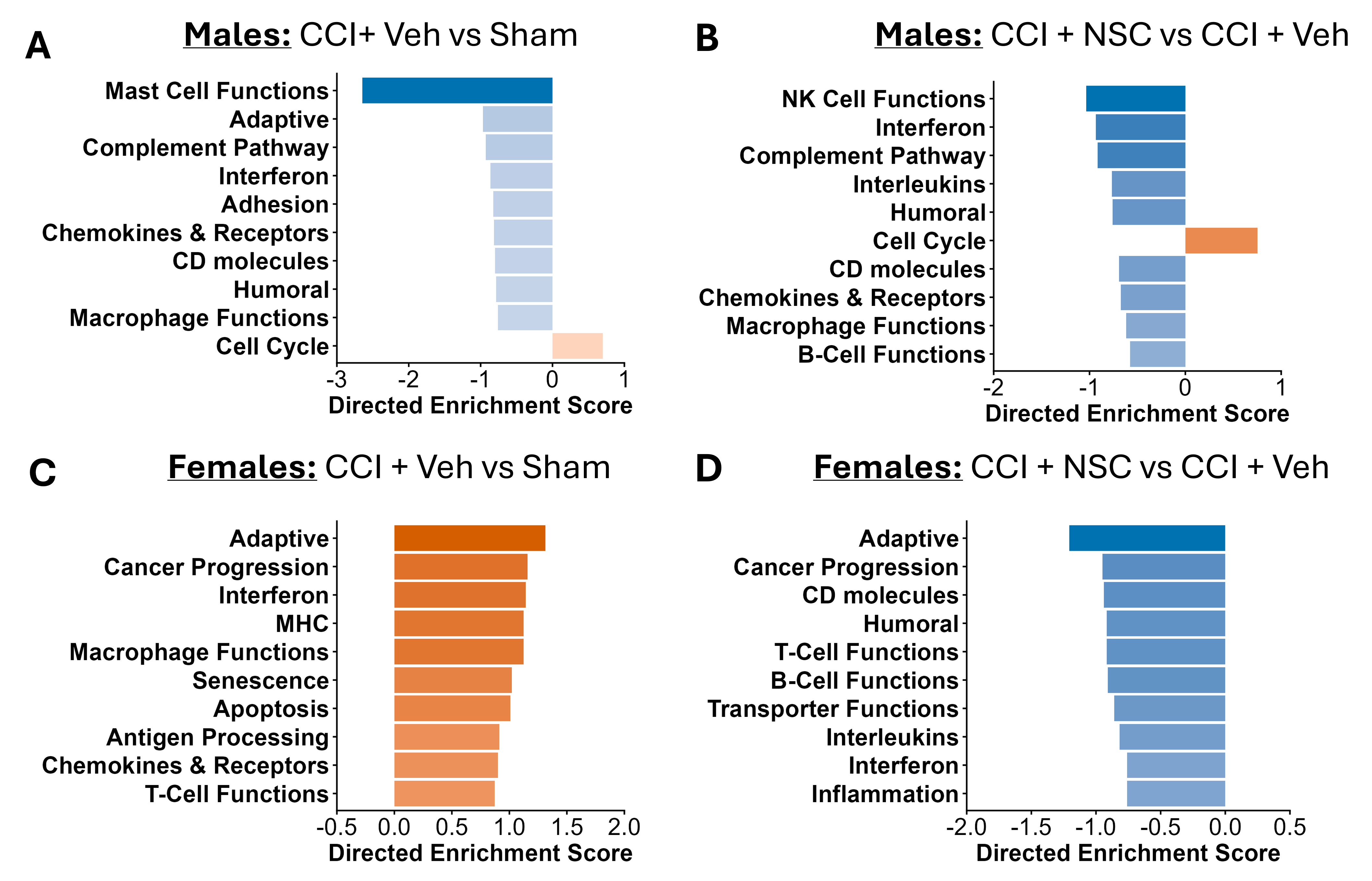

**Figure S3. NanoString functional pathway analysis of contralateral (CL) side tissue**

(A, B) Bar charts depicting functional enrichment analysis of gene expression in CL side (left hemisphere) brain tissue of male rats. The top ten significantly enriched pathways were identified using Gene Set Analysis in ROSALIND® (HyperScale architecture) with NanoString annotation databases for CCI + Veh vs Sham and CCI + NSC vs CCI + Veh comparisons. (C, D) Functional enrichment analysis of gene expression as in (A, B) in CL side (left hemisphere) brain tissue of female rats.

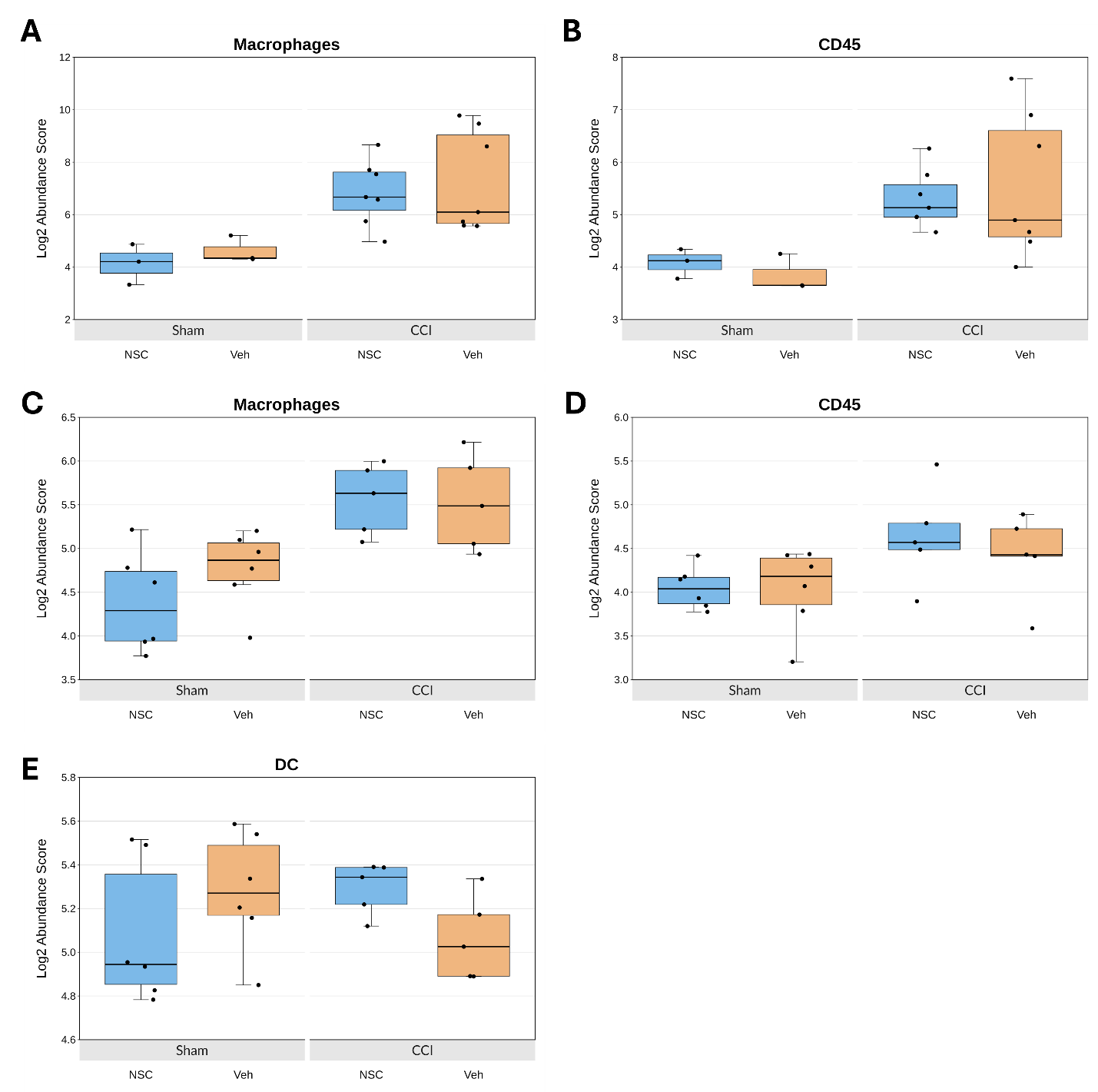

**Figure S4. Cell Type analysis.** NanoString Cell type profiling of Macrophages (A, C), CD45+ cells (B, D) and Dendritic cells (E) in Sham + NSC, Sham + Veh, CCI + NSC and CCI + Veh groups in female (A, B) and male (C, E) rats.

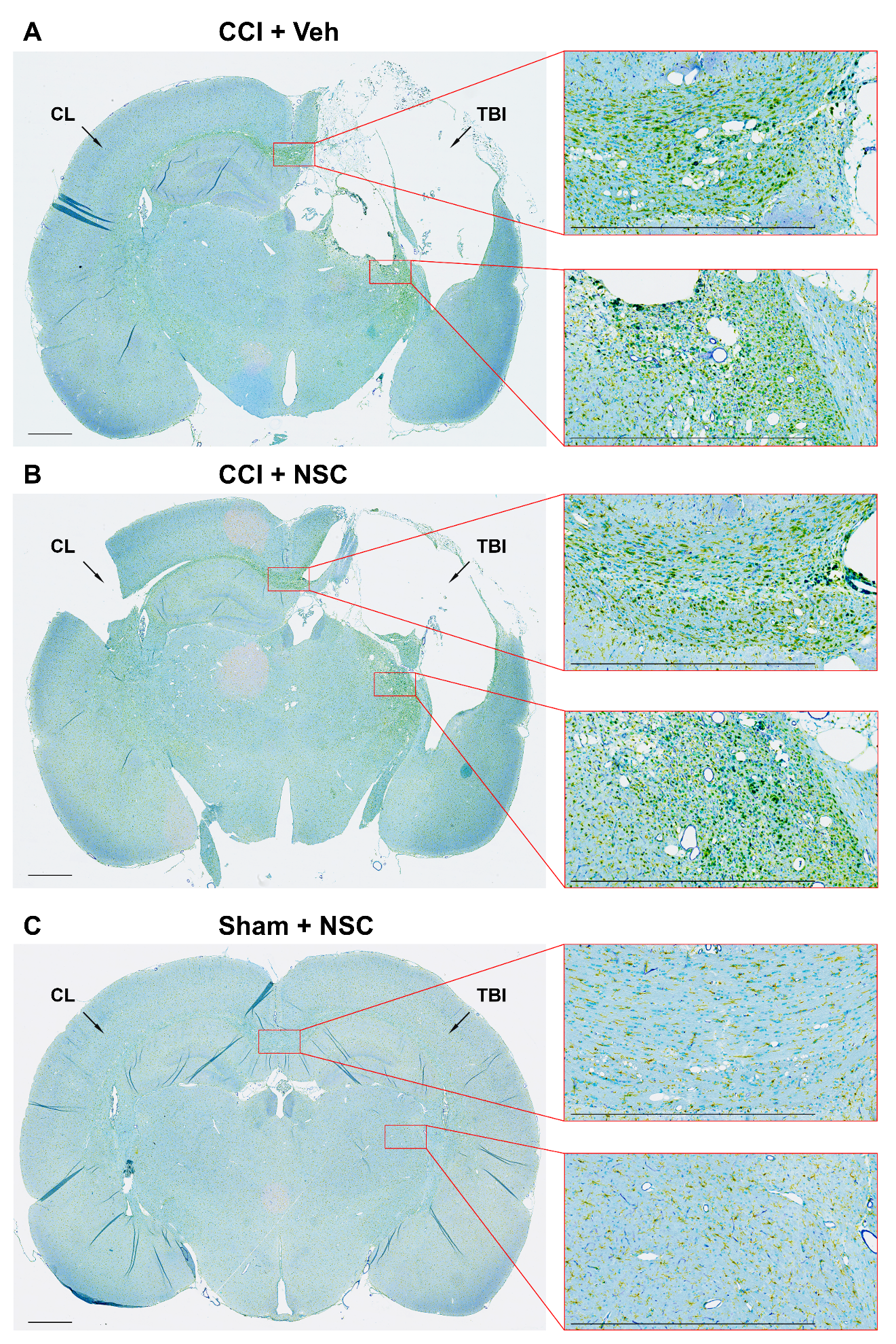

**Figure S5. Multiplex immunohistochemistry of rat brain tissue for IBA1 (Inflammation), CD68 (Immune Response), and CD31 (Vasculature).** IHC staining of TBI and CL hemispheres for IBA1 (yellow), CD31 (purple), and CD68 (teal) in brain tissue from Veh-treated CCI (A), LMNSC01-treated CCI (B), and Veh-treated Sham surgery rats (C). Areas of interest include the corpus callosum, and the thalamus on TBI side. Scale bars 1 mm.

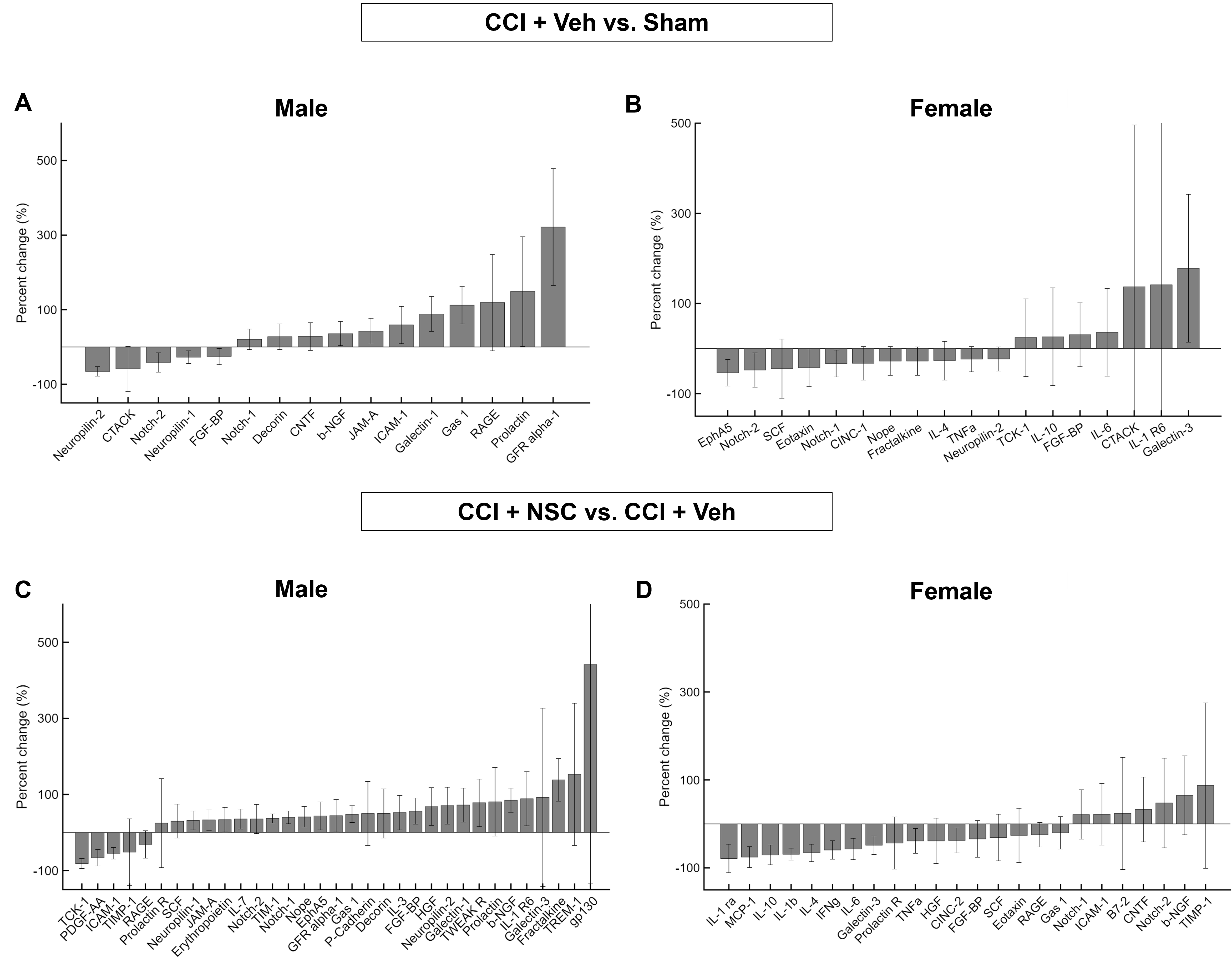

**Figure S6. Multiplex ELISA of CL side brain tissue protein expression after CCI surgery and LMNSC01 treatment.**

(A, B) Bar charts showing percent change in protein concentration in male and female rat brain lysates from CCI + Veh vs. Sham groups measured by multiplex ELISA in CL (left) side tissue.

Proteins with a percent change greater than ±20% are displayed. (C, D) Bar charts showing percent change in protein concentration of CL tissue from CCI + NSC vs. CCI + Veh groups.
